# Whole genome analysis reveals aneuploidies in early pregnancy loss in the horse

**DOI:** 10.1101/2020.02.25.964239

**Authors:** Charlotte A. Shilton, Anne Kahler, Brian W. Davis, James R. Crabtree, James Crowhurst, Andrew J. McGladdery, D. Claire Wathes, Terje Raudsepp, Amanda M. de Mestre

## Abstract

Chromosome abnormalities are well documented in human spontaneous abortion studies, yet rarely reported in domesticated animals. Rodent models have previously been used to study the effects of maternal ageing on oocyte quality and ultimately aneuploidy, however the differing endocrine profiles, oocyte characteristics and the polytocous nature of rodents are limitations for translation into human medicine. Early Pregnancy Loss (EPL) occurs in 5-10% of confirmed equine pregnancies and has no diagnosis in over 80% of cases. Aneuploidy has never been described in equine pregnancy loss, thus the objectives of this study were to quantify the frequency and characteristics of aneuploidy associated with equine EPL. EPL conceptuses were submitted from clinical cases of spontaneous pregnancy loss (14-65 days of gestation) between 2013 and 2018. Age matched control conceptuses were obtained from terminated clinically normal pregnancies (CNP). Aneuploidy was detected in 12/55 EPLs (21.8%), 0/10 CNP, 0/5 healthy term chorioallantois, and 0/5 healthy adult mares via genotyping. Whole genome sequencing (30X) and ddPCR validated results. Aneuploidies involved 10/32 equine chromosomes, consisting of nine trisomies and three monosomies. Autosomal aneuploidies were detected in both placental and fetal compartments in all samples tested. Aneuploid types (7/9) were mostly unique to EPL, supporting their embryonic/fetal lethality. Presenting the first evidence of aneuploidies in failed equine pregnancies not only provides the initial step in identifying genetic causes for these early losses, but also offers the horse as a new model for studying naturally occurring aneuploidy. We also demonstrate that SNP arrays provide a simple, cost effective way to screen aneuploidies across a large population.

**Author Summary:** The first 8 weeks of pregnancy is a critical time in both humans and horses, as the majority of pregnancy losses occur during this period. Despite such high prevalence, many cases do not have a known cause. Abnormal chromosome number (aneuploidy) is the most common finding in human pregnancy loss studies, but to date no equivalent study has been performed in domesticated animals, including the horse. We studied the genetics of naturally occurring pregnancy losses from Thoroughbred horses and found a similar level of aneuploidy to that observed in women. As humans and horses share similarities in their reproductive biology (ageing eggs, increased pregnancy loss in older mothers, similar key hormones), we suggest that by comparing the genetics of these two species, greater advances in identifying causes of aneuploidy pregnancy can be reached. Thoroughbred horses also tend to be more inbred than humans, facilitating the identification of mutations that increase the chance of aneuploidy, and this knowledge could potentially be applied in human medicine, as well as in species conservation.

## Introduction

Early pregnancy loss (EPL) represents the largest contributor to reproductive failure in many mammalian species [1–3]. In women, EPL causes emotional distress, and in domestic species it negatively impacts breeding economics, as well as introducing potential welfare issues [4, 5]. Between 5-10% of confirmed equine (*Equus caballus*) pregnancies end in the first 8 weeks [2] and despite intensive management strategies, the incidence has remained relatively stable over the past few decades [2, 6]. While there are many factors that increase a mare’s risk of EPL [7], these are not always causative *per se*, and in over 80% of EPLs no formal diagnosis is achieved [2], with the remaining 20% being diagnosed as either infectious (~4%) or non-infectious (~16%) in nature [2, 8]. In unexplained equine EPLs, genetic abnormalities are often attributed as a potential underlying cause, yet this remains unproven beyond a few case studies detailing balanced autosomal translocations in phenotypically normal mares with affected fertility [9–14]. Advancing maternal age has been consistently associated with pregnancy loss in many species [7, 15]. Whilst an abnormal uterine environment, more prevalent in aged mares, is likely to be a contributing factor, experiments involving the transfer of embryos from young mares into old mares (and vice versa) have suggested that this is not the main age-related underlying cause [16]. Oocytes from aged mares are of lower quality compared with younger fertile mares [17–19], indicating oocyte quality as a more likely risk factor for pregnancy loss.

Aneuploidy (gain or loss of a whole chromosome) is well documented in human spontaneous abortion [20, 21] and is associated with advancing maternal age. Some human autosomal trisomies (13, 18, and 21) have been identified in infants following live births, although these usually present with significant developmental pathologies [22] with only trisomy 21 individuals surviving to adulthood [23], and no documented surviving monosomies. The remainder of chromosomes that experience autosomal aneuploidy have only been identified in spontaneous abortion samples, leading to the theory that certain chromosomes do not tolerate aneuploidy and are therefore embryonic/fetal lethal.

While more commonly reported in humans, aneuploidy has also been noted occasionally in domesticated species including live born calves [24–26], dogs [27] and live born foals [28–30]. Equines reported to survive to term are often euthanised at a young age due to extreme developmental defects [28–30]. In species with reported live born individuals with autosomal aneuploidies, all are trisomies with no reported monosomies in the literature [31]. This indicates that while some trisomies may be tolerated to term, monosomies are always lethal at some stage during pregnancy. Very few studies have investigated aneuploidy rates in spontaneous abortion of any non-human species. One study karyotyped 55 bovine spontaneous abortions [32] identifying 6 individuals with chromosomal abnormalities (4 trisomies, 1 translocation, and 1 male/female chimera). This has led to the suggestion that high rates of aneuploidy associated with pregnancy loss might be a human-specific event. Investigations analysing *in vitro* or *in vivo* generated blastocysts have identified chromosomal abnormalities in a number of species including equine [33], bovine [34], ovine [34], and leporidae [35]. It is important to note that these blastocysts have been purposely prevented from achieving a pregnancy, and therefore the outcome of these chromosomal abnormalities cannot be determined. Further to this, the relevance of these observations in blastocysts to naturally conceived pregnancies is untested.

Previous attempts to identify chromosomal abnormalities in equine abortions have been largely unsuccessful [36, 37], but the successful isolation of conceptus material and culture of trophoblast cells from early abortus material [38] opens up the possibility for investigation into genetic and chromosomal causes of EPL in the mare. While karyotyping of cells has been used as the gold standard for aneuploidy analysis, it requires much expertise and time to generate results in large numbers. As technologies have progressed, new methods are emerging that may replace karyotyping as gold standard, allowing for higher throughput analysis and diagnosis of aneuploidy, even in degrading conceptus tissues. Whole genome sequencing (WGS) is one potential method, however the cost is still too high for this to be a viable solution. Single Nucleotide Polymorphism (SNP) Arrays are a lower cost, high throughput method that may offer an answer. A low-density SNP array (Equine SNP50 BeadChip) has previously been used to detect aneuploidy in two live born horses suspected of chromosomal abnormalities [30], offering evidence for the validity of this methodology.

In order to explore whether aneuploidy is a feature of failed pregnancies in domesticated animal species, we utilised methods previously reported [38] to generate a large bank of conceptuses from naturally occurring clinical cases of EPL in mares. We hypothesised that due to the rarity of aneuploidy in foals born at term, aneuploidy presents as embryonic/fetal lethal and will be detectable in both placental and fetal compartments consistent with possible origins in maternal meiosis. The primary aim of this study was to quantify the frequency and characteristics of aneuploidy associated with EPL in the mare.

## Results

### Descriptive data of SNP array sample population

The median gestational age of the early pregnancy loss (EPL; n=55) and clinically normal pregnancies (CNP; n=10) conceptuses was 42(±12.5SD) and 33.5(±10.1SD) days respectively (Fig 1A). The range of gestational ages was 14-67 and 29-64 days for EPL and CNP respectively and were not significantly different (p=0.1957) (Fig 1A). There was no significant differences in the median maternal age of the EPL and CNP samples (10±4.9SD and 9.5±6.3SD years respectively, p=0.8967), which ranged between 3-12(EPL) and 2-20(CNP) years (Fig 1B). All EPL conceptuses came from individual mares, while the CNP conceptuses came from a pool of 7 mares, 3 of whom provided two conceptuses. Male and female conceptuses were equally represented on the array (p=0.167) (Fig 1C). Chorioallantois tissue was obtained from five term births following the normal delivery of a healthy foal. DNA from the dams of the CNPs were used as healthy adult controls. The five Thoroughbred mares aged 2 to 20 years were reproductively sound.

**Fig 1.**
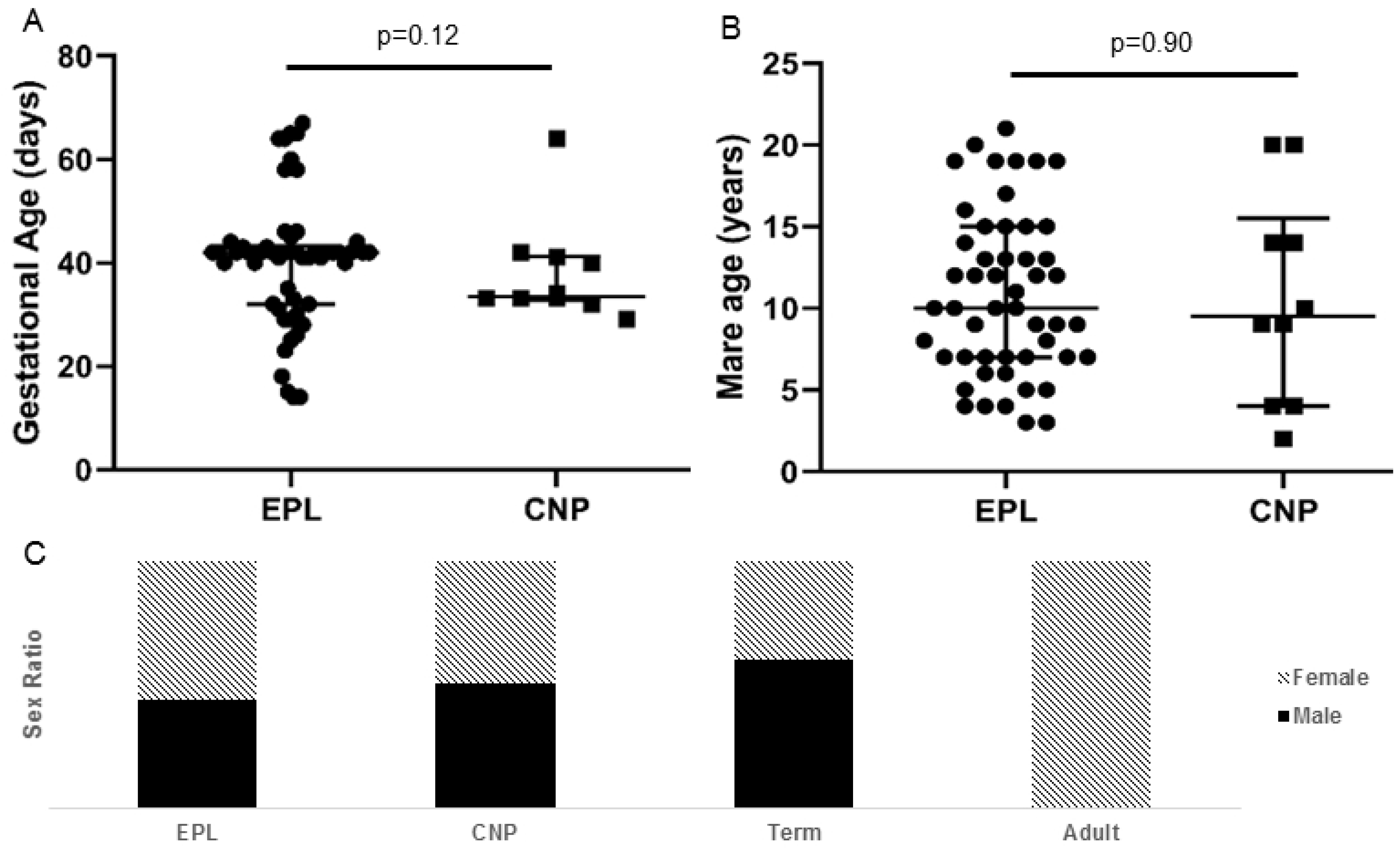
Description of sample population on the SNP Array. No significant difference was found in the average (A) gestational age or (B) mare age of pregnancies within the early pregnancy loss (EPL) and clinically normal pregnancy (CNP) groups. Mean with standard deviation plotted. (C) Males and females were equally represented on the array for the EPL, CNP, and healthy term placentae. All adults were females.

### Concordance of SNP genotypes

Concordance analysis of the SNP genotype calling was used to assess the accuracy/repeatability of the SNP array (Fig 2A). Matching allantochorion (ALC) and fetal samples (from the same conceptus) had a median concordance of 98.7% (range 97.4-99.3) and were significantly different to samples not known to be related (median concordance of 70.7%, range 62.9-78.6, p<0.0001). Samples from pregnancies known to be half siblings were not significantly different from those known to be full siblings (median concordance 74.7 and 81.2, respectively, range 67.0-76.7 and 81.1-83.0, respectively). Tissue repeats (different regions of the same allantochorion) had a median concordance of 99.1% (range 98.9-99.4) while technical replicates (different aliquots of the same DNA sample) had a median concordance of 99.1% (range 99.0-99.4). The sex of the conceptus was determined by PCR (Fig 2B) and compared with the predicted sex based on the copy number of X chromosome as visualized by IGV (Fig 2C). There was 100% agreement (75/75 individuals tested) in the sex of the tissue as determined by PCR and IGV visualisation.

**Fig 2.**
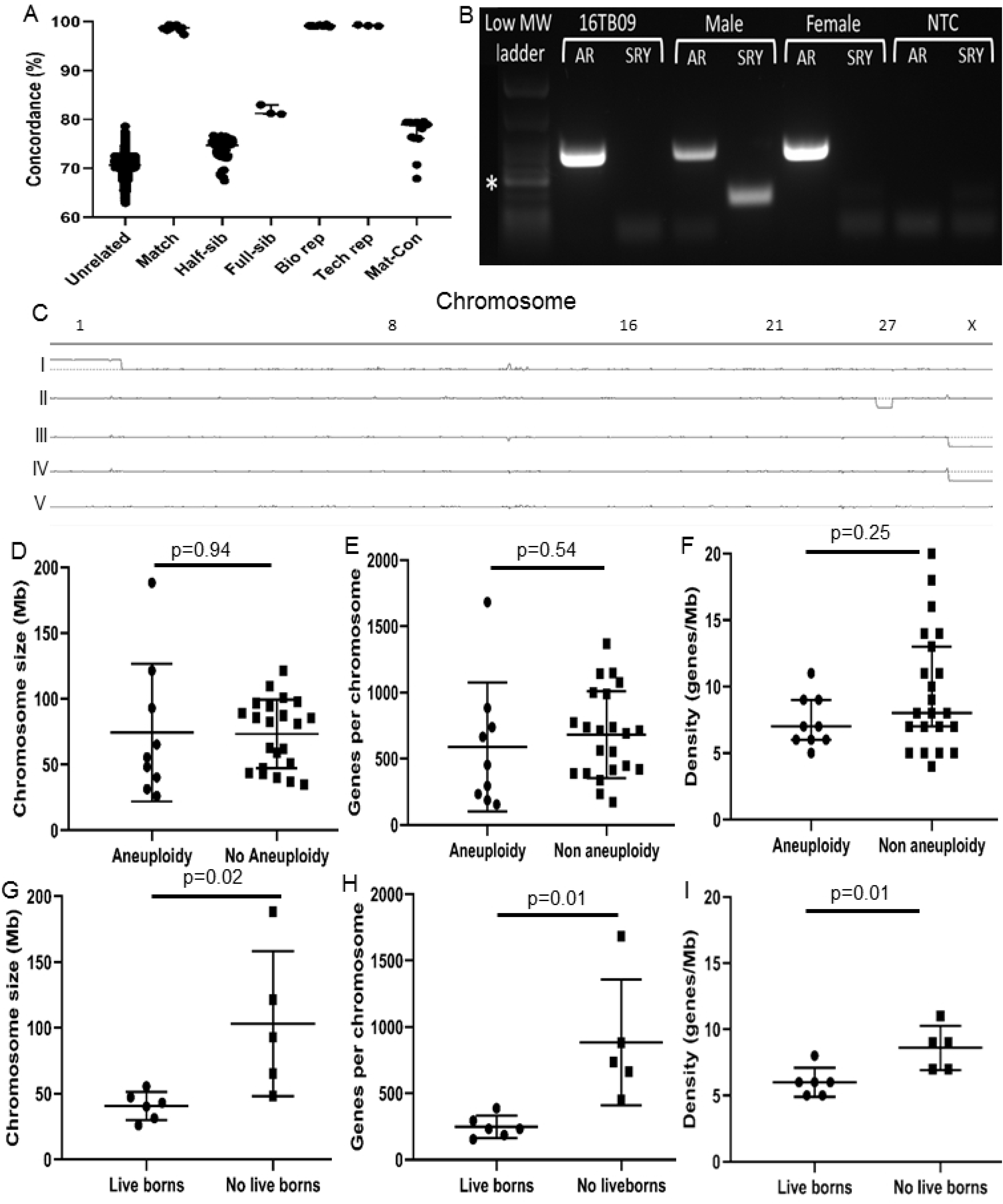
Initial validation of results and identification of aneuploid chromosomes. (A) Concordance analysis of the SNP calls for each sample (n=96). Samples from the same conceptus (“Match” – matching fetal and allantochorion DNA) were significantly different to samples that were not known to be related at all (“Unrelated”, p<0.0001). “Half-sib” compared the concordance of individuals sharing one parent, while “Full-sib” compared those that shared both parents. “Bio rep” indicates the comparison of three regions across the allantochorion of a single conceptus, while “Tech rep” compared a single aliquot of DNA from the same region of allantochorion of a single individual. “Mat-Con” compared the concordance of maternal DNA with the corresponding offspring DNA. Median with interquartile range plotted. (B) Sex determination using standard PCR with primers for Sex determining Region of Y (*SRY;* Y chromosome; 131 bp) and Androgen Receptor (*AR*; X chromosome; 293 bp) validated the X chromosome copy number status. Confirmed male and female equines as positive controls, and ddH_2_O as no template control (NTC). *200bp band on low MW ladder. (C) Examples of whole genome copy number visualisation with Integrative Genomics Viewer. Chromosome number is displayed horizontally across the top axis, with the centre horizontal line indicating a copy number of 2 (diploid). Allantochorion of (I) female trisomy 1 EPL, (II) female monosomy 27 EPL, and (III) male diploid CNP, along with (IV) male diploid term chorioallantois and (V) female adult peripheral blood mononuclear cells. (D-I) Analysis of chromosome characteristics comparing D) and G) chromosome length, E) and H) the total number of genes, and F) and I) the gene density per chromosome. Top panel compares the autosomal chromosomes that were found to be aneuploid within the EPL subpopulation of this study to those not identified as involved in aneuploidy (n=31 for each graph). Bottom panel compares characteristics of aneuploid autosomal chromosomes previously reported in live born equines with those unique EPLs in this study (n=10 per graph). Mean with standard deviation plotted.

### Aneuploidies identified in failed equine pregnancies

Aneuploidy of at least one chromosome was found in 12/55 EPL pregnancies (21.8%), as compared with 0/10 CPN manually terminated pregnancies, 0/5 healthy term, and 0/5 healthy reproductively sound adult mares (Fig 2C, Table 1). Aneuploidies were noted on 10/32 chromosomes (1, 3, 15, 20, 23, 24, 27, 30, 31, and X), representing both trisomies (9/12) and monosomies (3/12). Aneuploidies were noted as single events in each pregnancy (only one chromosomal imbalance event), except for chromosomes 23 and 24 which occurred together in a single failed pregnancy (Table 1). Only 2 of the autosomal aneuploidy types have previously been reported in live born equines (trisomy 30 and trisomy 23) with the remaining 7 aneuploidy types being unique to this study (Table 2).

**Table 1.** Aneuploidy noted in 12/55 early pregnancy loss (EPL) equine conceptuses.

| Sample ID | Sex | Gestational Age (days) | Maternal Age (years) | Maternal Breed | Aneuploidy noted |
| --- | --- | --- | --- | --- | --- |
| 14AI01 # | F | 14 | 4 | WB | Trisomy 1 |
| 18TB10 | M | 31 | 6 | TB | Trisomy 3 |
| 18TB07* | F | 28 | 19 | TB | Trisomy 15 |
| 14TB02 | M | 42 | 19 | TB | Trisomy 20 |
| 16TB02 | F | 58 | 13 | TB | Trisomy 20 |
| 18TB09* | M | 42 | 3 | TB | Trisomy 23 and 24 |
| 16TB09* | F | 60 | 10 | TB | Monosomy 27 |
| 18TB08 | F | 42 | 19 | TB | Monosomy 27 |
| 16TB03 | M | 32 | 19 | TB | Trisomy 30 |
| 18TB05* | F | 41 | 9 | TB | Trisomy 30 |
| 18TB15 | F | 29 | 10 | WB | Monosomy 31 |
| 17TB02 | F | 26 | 13 | TB | Trisomy X |

All samples from allantochorion (ALC) except #yolk sac of day 14 conceptus. \*conceptuses with both ALC and fetal (F) tissue present on the array, F = female, M = male, TB = Thoroughbred, WB = Warmblood

**Table 2.** Comparison of autosomal aneuploidy types unique to our study (rows above the black line) and those also reported in live born equines (below the line).

| Aneuploidy type | Chromosome size (Mb) | Genes per chromosome (n) | EPL conceptuses n | Maternal age (years) | Live born n | Maternal age (years) |
| --- | --- | --- | --- | --- | --- | --- |
| Trisomy 1 | 188.3 | 1683 | 1 | 4 | 0 | NA |
| Trisomy 3 | 121.4 | 883 | 1 | 6 | 0 | NA |
| Trisomy 15 | 92.9 | 664 | 1 | 19 | 0 | NA |
| Trisomy 20 | 65.3 | 738 | 2 | 13 and 19 | 0 | NA |
| Trisomy 24 | 48.4 | 453 | 1* | 3 | 0 | NA |
| Monosomy 27 | 40.3 | 232 | 2 | 10 and 19 | 0 | NA |
| Monosomy 31 | 26 | 154 | 1 | 10 | 0 | NA |
| Trisomy 23 | 55.6 | 294 | 1* | 3 | 1 | –<sup>a</sup> |
| Trisomy 26 | 43.1 | 232 | 0 | NA | 1 | 3<sup>f</sup> |
| Trisomy 27 | 40.3 | 232 | 0 | NA | 3 | 5<sup>b</sup>, 26<sup>c</sup>, –<sup>d</sup> |
| Trisomy 28 | 47.3 | 388 | 0 | NA | 1 | 14<sup>e</sup> |
| Trisomy 30 | 31.4 | 185 | 2 | 9 and 19 | 2 | 23<sup>f</sup>, –<sup>d</sup> |
| Trisomy 31 | 26 | 154 | 0 | NA | 1 | 26<sup>g</sup> |

EPL = early pregnancy loss, NA = not applicable, \* = same conceptus trisomic for two different chromosomes, – = unreported mare age. <sup>a</sup> [39], <sup>b</sup> [29], <sup>c</sup> [28], <sup>d</sup> [30], <sup>e</sup> [40], <sup>f</sup> [41], <sup>g</sup> [42].

Aneuploidy was detected in both large and small autosomes (range 26-188.3 Mb), along with chromosome X. There was no significant difference between the mean length (±SD) of the autosomal chromosomes detected as aneuploidy compared with those undetected (74.4±52.4 and 73.3±26.0 Mb respectively, p=0.25) (Fig 2D). The mean number of genes per autosomal chromosome did not significantly differ between the aneuploidy and non-aneuploidy chromosomes (587.3±486.9 and 681.5±326.9 respectively, p=0.53) (Fig 2E). The median density of autosomal chromosomes did not significantly differ between those detected as aneuploidy and those not detected (8±4.5 and 7±1.9 genes per Mb, respectively, p=0.246) (Fig 2F).

The mean length (±SD) of autosomal chromosomes previously reported in aneuploidy live born foals (40.6 Mb ±10.71) were significantly shorter than those that are unique to this study and only reported in EPL conceptuses (103.3 Mb ± 55.04) (p=0.022) (Fig 2G). The mean number of genes on each autosomal chromosome was significantly different between chromosomes reported in live born and those unique to this study (247.5±83.70 and 884.2±472.7 genes per chromosome, respectively, p=0.01) (Fig 2H). Autosomal chromosomes not previously documented in aneuploidy individuals had on average a significantly higher gene density compared with those that have also been reported in the literature (6±1.1 and 8.6±1.7 genes per Mb, respectively, p=0.013) (Fig 2I).

### Validation of SNP Array results by WGS and ddPCR

Whole genome sequencing (WGS) was performed on 5 CNP, 6 EPL diploid conceptuses, and 1 EPL (18TB08) with monosomy 27 according to SNP genotyping. Average read coverage for chromosome 27 in 18TB08, was approximately half of that for other chromosomes, indicating a monosomy (Fig 3A). For all other WGS samples without aneuploidy indicated by SNP array analysis (n=11), the average read coverage for each chromosome was equal (Fig 3A). Next, digital droplet polymerase chain reaction (ddPCR) was performed to further validate the presence of trisomy 1 (14AI01) (Fig 3B) and monosomy 27 (16TB09 and 18TB08) (Fig 3C) using genes on chromosome 18 as a reference. Two control samples (CNP; 1806, 1808) had an approximate copy number of 2 for both genes on chromosome 1 (*ACTC1* and *SHTN1*). The trisomy 1 EPL had an approximate copy number of 3.5 for both genes on chromosome 1 while the two monosomy 27 EPLs that were diploid for chromosome 1 had an approximate copy number of 2. The CNPs (1806, 1808) and the trisomy 1 EPL had an approximate copy number of 2 for both chromosome 27 genes *NRG1* and *ANGPT2* while the two monosomy 27 EPLs had an approximate copy number of 1 for both genes (Fig 3C).

**Figure 3.**
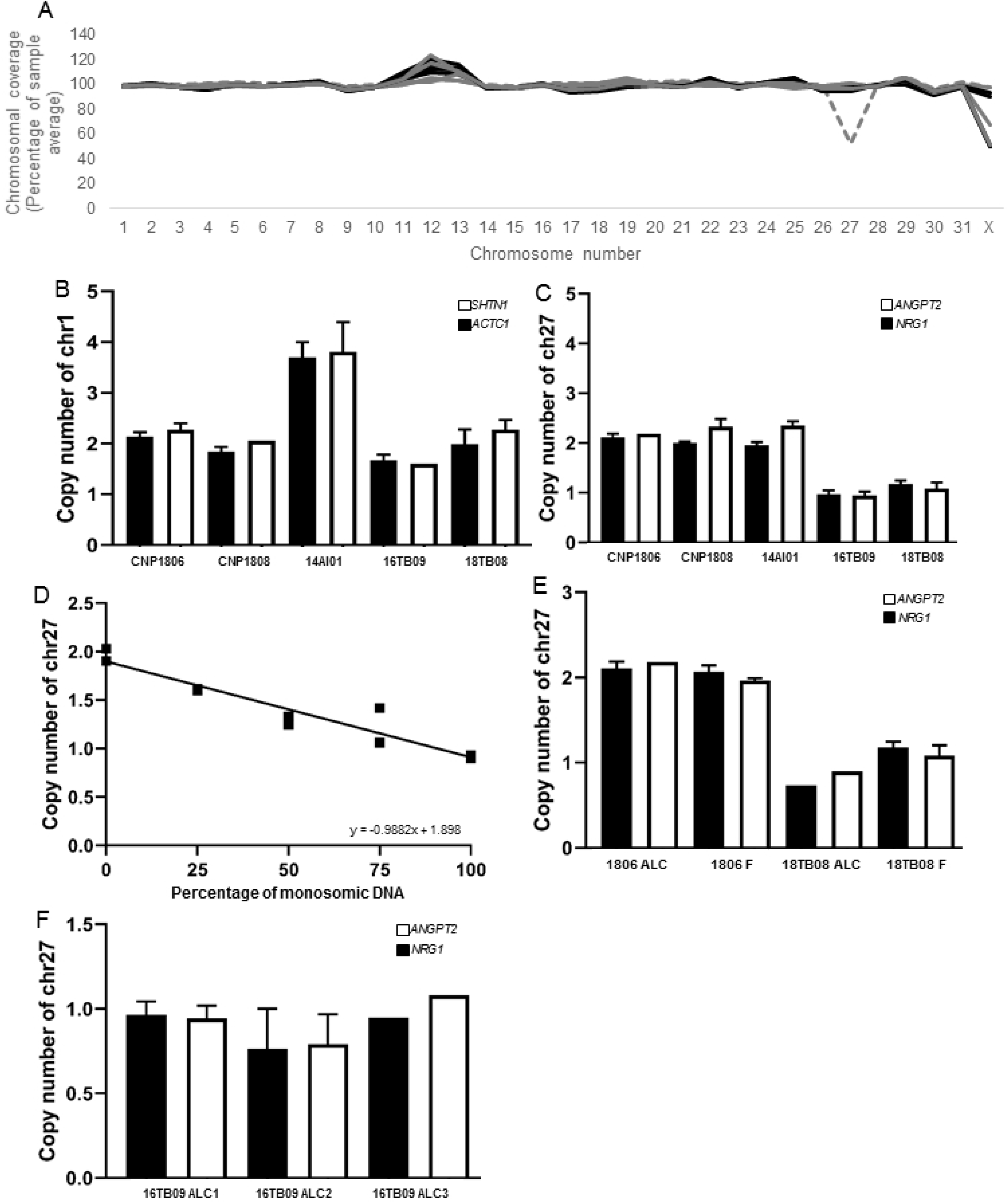
Validation of results by Whole Genome Sequencing and digital droplet PCR. (A) Whole genome sequencing (WGS) of 12 of the array samples (n=1 aneuploidy EPL grey dashed line, n=3 non aneuploidy EPL grey, n=4 CNP black). Average coverage per chromosome calculated with SAMtools from GATK. Graph presents the chromosome as a percentage of the autosomal average. Digital droplet PCR (ddPCR) of (B) chromosome 1 genes *ACTC1* and *SHTN1*, and (C) chromosome 27 genes *NRG1* and *ANGPT2* across 5 samples (n=2 CNP, n=1 trisomy 1 EPL, n=2 monosomy 27) relative to the reference region (*MCM6*) on chromosome 18. Primers were designed for two regions at the end of each chromosome. All samples were analysed in duplicate. (D) ddPCR of chromosome 27 (*NRG1*) in duplicate. Diploid and monosomy 27 DNA was mixed at different ratios to represent varying levels of mosaicism. Copy number was normalised to the *MCM6* reference region on chromosome 18 and all samples were analysed in duplicate. Negative correlation was noted between the copy number of chromosome 27 and increasing concentration of monosomic DNA (R = −0.9882, p<0.0001). (E) DNA from allantochorion (ALC) and fetus (F) of two different conceptuses (n=1 diploid CNP, n=1 monosomy 27 EPL), analysed in duplicate for the two regions of chromosome 27 genes. (F) DNA from three different regions of allantochorion (ALC) of 16TB09 (monosomy 27) analysed with ddPCR to identify whether conceptus 16TB09 was a mosaic. All regions were analysed in duplicate with chromosome 27 genes and normalised to the *MCM6* reference on chromosome 18. Error bars indicate standard deviation.

### Mosaicism is not a common feature of equine aneuploid conceptuses

Aneuploid human pregnancies have been shown in some cases to be mosaic [43, 44] so we investigated the ploidy status within and across fetal tissues. DNA sequences of matching allantochorion (ALC) and fetal (F) tissues from 14 pregnancies (n=8 EPL, n=6 CNP) were assessed using IGV. Where aneuploidies were identified in the ALC gDNA (16TB09, 18TB05, 18TB07, 18TB09) the same aneuploidies were always noted in the fetal gDNA. Matching ALC and F tissues of non-aneuploid conceptuses, also shared the same diploid status (n=10). An artificial mosaic (varying ratios of normal and aneuploid DNA) was analysed by ddPCR and demonstrated what a mosaic monosomy result would look like when analysed with ddPCR. The aliquot with 100% diploid DNA had a copy number of 2 for chromosome 27, while the aliquot with 100% monosomic DNA had a copy number of 1 for chromosome 27. The copy number of chromosome 27 was negatively correlated (R = −0.9882, p<0.0001) with the increasing volume of monosomic DNA (Fig 3D). Digital droplet PCR (ddPCR) for two genes on chromosome 27 (*NRG1* and *ANGPT2*) further confirmed the diploid status of both ALC and F from a CNP conceptus (1808), and the monosomy 27 status of both ALC and F from an EPL (18TB08) (Fig 3E). Next, we investigated the presence of monosomy 27 in three independent ALC tissue samples collected from different regions of the placentae. All three regions of the ALC tested were found to have monosomy 27 (Fig 3F).

### Phenotypes of EPL and clinical analysis

Of the 55 EPL pregnancies analysed, 34 had a fetus confirmed present within the conceptus. There was no significant difference in aneuploidy occurrence between conceptuses presenting with a fetus and those without a fetus (p=0.361). Of the 12 aneuploidy conceptuses, 3 did not have a fetus at dissection; one was too young for an embryo proper (Trisomy 1, 14 days old), one was submitted as a complete and intact conceptus with a suspected embryonic disc only with no evidence of vasculature (Monosomy 31, Fig 4A – age matched CNP for comparison Fig 4B), and one had evidence of a fetus recorded in clinical records but no fetus was present at dissection (Trisomy X). One submission was accompanied by autolytic fetal remnants (trisomy 20) and the remaining 7 aneuploid EPLs presented with an intact/partially intact fetus of variable phenotypes. A 32 day trisomy 30 embryo proper (Fig 4C) appeared to have distorted and mismatched developmental features compared with the age matched CNP embryo proper (Fig 4D), although autolytic changes impaired full assessment of this EPL specimen. The day 60 monosomy 27 fetus (Fig 4E) appeared oedematous and congested when compared with the day 64 CNP fetus (Fig 4F) consistent with an abnormal vasculature phenotype.

**Figure 4.**
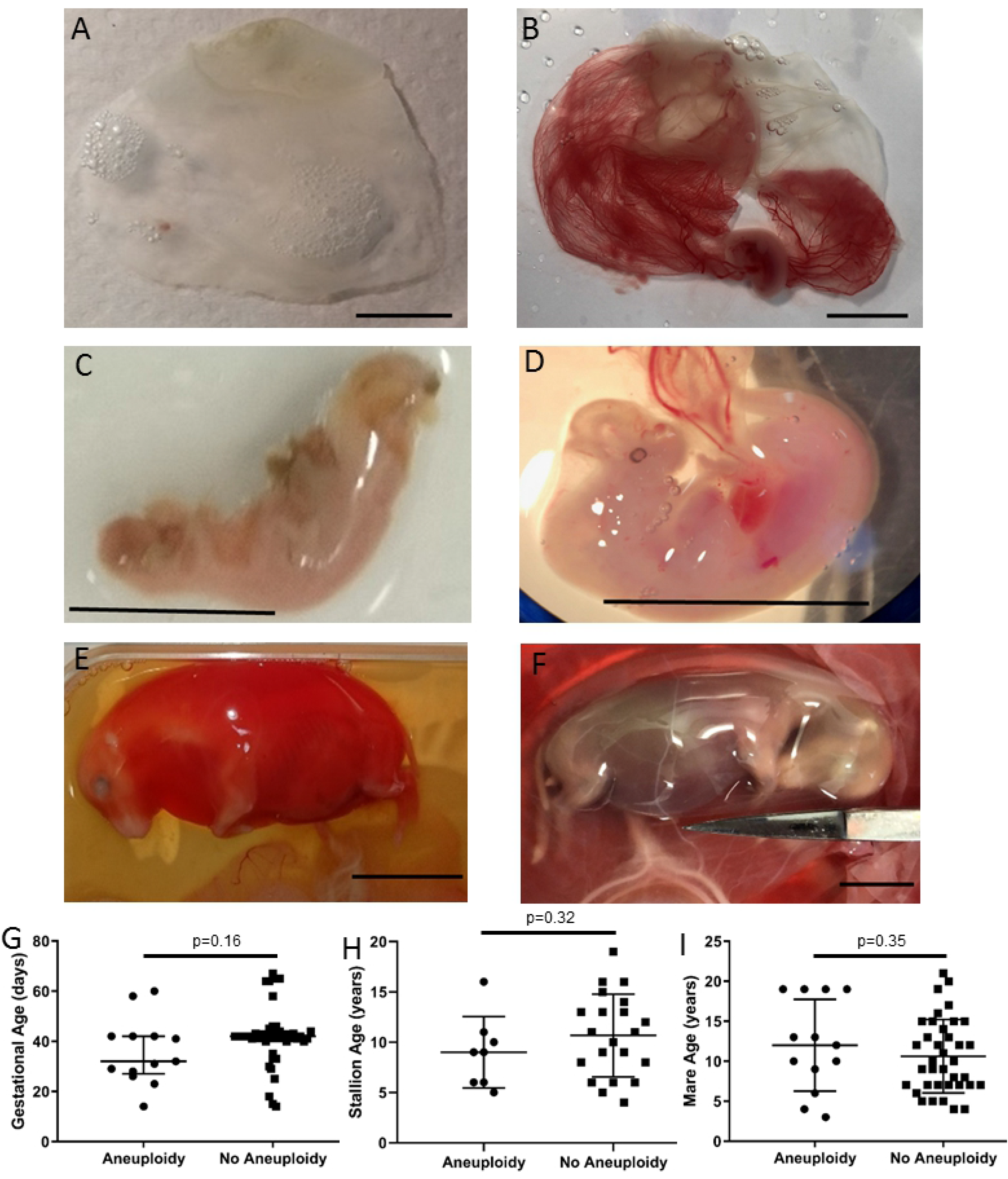
Phenotypes and clinical analysis of aneuploidies. (A) Monosomy 31, 29 day gestation failed conceptus and (B) age matched 29 day clinically normal conceptus. (C) Trisomy 30, 32 day gestation failed embryo proper and (D) age matched 33 day clinically normal embryo proper. (E) Monosomy 27, 60 day gestation failed fetus and (F) age matched 64 day clinically normal fetus. Scale bar = 1cm for all images. (G) Gestational age, (H) stallion age, and (I) mare age did not significantly differ between aneuploidy EPLs and non-aneuploidy EPLs. Mean with standard deviation plotted.

None of the variables analysed (conceptus sex, maternal age, paternal age, gestational age, breed, use of ovulation induction, twin pregnancies, positive bacterial growth from a uterine swab at time of loss) were significantly associated with risk of aneuploid EPLs (Fig.4G-I, S1 Table). Next, we compared the likelihood of a mare ending the season with a live foal after suffering either an aneuploid EPL or a non-aneuploid EPL. Mares who suffered an EPL after 43 days when endometrial cups are established were excluded. Mares who suffered aneuploidy EPL were significantly more likely to end the season with a live birth (3/4 mares) compared to those with a diploid EPL (2/15 mares) (p=0.0374).

## Discussion

An abundance of reports into human miscarriages, cite aneuploidy as the greatest cause of the pregnancy failure [20, 21]. In contrast, reports of aneuploidy associated with pregnancy loss in domesticated animals is rarely reported, and never in the horse. Here we identified autosomal aneuploidies in 20% of naturally occurring equine EPL conceptuses, slightly lower than the 35-50% reported in women [20, 45] possibly explained by the shorter gestational time frame used in our study. Seven of the aneuploidy types we described were unique to the study, including two monosomies. Further, none of the pregnancies assessed across both fetal and placental compartments were mosaic, collectively providing evidence that these aneuploidies are likely to be embryonic/fetal lethal and zygotic in origin. Additionally, the Axiom™ Equine Genotyping Array proved to be a successful methodology to identify aneuploidies verifiable by whole genome sequencing and ddPCR.

In mammalian species studied to date, very few aneuploidies are tolerated to term. Whereas partial monosomies have been identified in humans [46], no cases of complete autosomal monosomy compatible with birth have been recorded in any species. Therefore, it is highly likely the monosomic conceptuses described here involving chromosomes 27 and 31 failed due to decreased gene dosage resulting in embryonic/fetal lethality. While trisomies can present in live borns, the best known being trisomy 13 and 21 in humans, the phenotypes vary considerably [47]. In humans, the phenotypes of trisomy 21 (Down’s syndrome) include fetal loss [48], heart disease [49] and early onset Alzheimer’s disease [50]. In the horse, there are only 8 individual case reports of aneuploid live borns, involving chromosomes 23, 26, 27, 28, 30, and 31. These presented with variable congenital abnormalities involving the musculoskeletal, neurological, and vasculature systems [28–30, 39–42]. Of these six equine aneuploidy types tolerated to term, only trisomy 30 was also identified in EPL conceptuses. How equine trisomy 30 can result in both a fetal lethal and live born phenotype is not known, although studies of similar phenotypic variation in Down’s Syndrome are reported, suggesting the elevated transcript levels generated by the genotype could be modified by genes on other chromosomes and/or the environment to initiate and exacerbate the resultant phenotype [51].

The remainder of the trisomy EPLs identified here have not previously been reported. The aneuploid chromosomes unique to failed pregnancies were found to be on average significantly larger, more dense, and with more genes compared with those that present a mixed phenotype (EPL/live born) consistent with the hypothesis that duplication or deletion of larger chromosomes are more likely to result in greater genetic imbalances and hence earlier lethality. Indeed, amongst our autosomal trisomies, those involving the largest chromosomes (trisomy 1, 3, and 15) were from pregnancies that failed at the youngest gestational ages (days 14, 32, and 28, respectively). The equine pregnancies assessed here were all clinically recognised, therefore the overall incidence of aneuploidy in equine pregnancy is predicted to be higher than the 22% we report. In support of these, there is a report of aneuploidy of larger chromosomes (2 and 4) in *in vitro* and *in vivo* generated equine blastocysts [33]. These aneuploid types were not identified here but given the size of these two chromosomes, it is plausible that their phenotype is embryonic lethal prior to clinical detection.

Mosaicism indicates a somatic mutation that occurred during the mitotic divisions of embryo development, with a negative correlation between developmental time at error and the degree of mosaicism. The combination of aneuploidy type, the degree of mosaicism, and the tissue compartment location determines the severity of the phenotype [52]. Mosaicism, and the percentage of the cells that it inhabits, offers an insight into the initial starting point of the aneuploidy. For a true aneuploidy (100% of cells containing the imbalance), the imbalance must have occurred prior to conception. The imbalance more commonly occurs within the oocyte (during meiosis I or II), with as high as 93% of human trisomy 21 cases being maternal in origin, but can also occur within spermatocytes [53]. Here we tested the matching allantochorion (ALC) and fetus (F) of 9 EPL and 6 CNP conceptuses, and multiple allantochorion samples from two separate conceptuses using a combination of SNP analysis and ddPCR. We found that the aneuploidy status was consistent between and within tissue compartments in all individual pregnancies tested, indicating a true aneuploidy instead of a mosaic and suggesting they were maternally or paternally derived, as is commonly found in humans [53].

Advancing maternal age is associated with decreased quality of oocytes (reviewed in [54]), from altered epigenetic profiles [55] and mitochondrial DNA deletions [56]. Aged oocytes also display altered microtubule spindle alignment [57] and weakened centromere cohesion [58] resulting in increased risk of aneuploidy. In the mare, advancing maternal age significantly increases the risk of chromosome misalignment [19] and early pregnancy loss [7]. Therefore we proposed that advancing mare age would increase the risk of aneuploidy. Whilst numerically there was an increase in the proportion of aneuploid pregnancies in older mares, there was also an increase in the proportion of very young mares with aneuploid pregnancies and neither were statistically different to the non-aneuploid EPL pregnancies. It is possible that this reflects the fact that non-aneuploid EPLs are also at increased risk with advanced maternal age, as age is associated with increased endometrial disease [59]. To accurately assess the impact of maternal age on aneuploidy, we would need to compare the maternal age of aneuploidy pregnancies to a cohort of mares with successful pregnancies over the same years, but due to the spread of mares across multiple practices, this was not realistically achievable within this study. The identification of embryonic lethal aneuploidy EPLs in very young mothers has been reported in human medicine [60].

Advancing paternal age at the time of conception has been linked with an increased risk of congenital abnormalities in humans [61] due to the increased risk of mitotic errors associated with continuous spermatogenesis, with approximately 1-2% of spermatozoa from donors being aneuploid [62]. It has been investigated as a potential risk factor for aneuploidy in human pregnancy losses, with no significant associations detected [63]. We also found that paternal age has no association with an increased risk for aneuploidy EPLs, however paternal age may still influence the risk of smaller genetic abnormalities.

While mice models of oocyte ageing has led to evidence of DNA fragmentation [64] and an altered gene expression patterns [65], the fundamental differences between rodents and humans may mean that these findings are not always transferrable to human medicine. Rodents are polytocous and display differing endocrine profiles compared with humans. Mares, on the other hand display similar endocrine profiles (notably follicle stimulating hormone levels) to women, particularly with regards to ageing (reviewed in [66]). As discussed above, advanced maternal age is associated with higher incidence of pregnancy loss in both species [2, 15]. Our study highlights that aneuploidy rates are similar between human and horse natural occurring pregnancy losses, show a similar propensity to be maternally or paternally derived, and a possible bias towards very young and old mothers being at risk. We suggest that the horse may be a viable alternative system to study the underlying mechanisms of aneuploidy and in the future test novel therapeutics, for application in human medicine.

Of the 20 individuals with autosomal aneuploidy reported to date (combining reports in the literature with our study), it is of note that approximately 45% involve chromosome 27 or 30 (5 trisomy live born [28–30, 41], two monosomy 27 EPLs, and 2 trisomy 30 EPLs). The underlying mechanisms of why certain chromosomes would be more susceptible to aneuploidy is not known with further studies warranted. Chromosomes 27 and 30 are among the most gene poor equine autosomes (6 genes/Mb each), possibly explaining their higher incidence. The difference in lethality of chromosome 27 is interesting as it appears that trisomies of this chromosome are viable to term, and display varying characteristics within 24 months of birth including cryptorchidism, bilateral carpal flexural deformity and an inability to suck [28], skeletal abnormalities and a failure to thrive [30], and atypical gait and reduced social skills [29]. Our study has added to the list of chromosome 27 aneuploidies by documenting 2 monosomies in early failed pregnancies. Both of these monosomies displayed extreme oedema with intact vasculature. The oedematous and congested phenotype of one of the EPLs indicates a potential for cardiac failure. The trisomy 30 live borns included a small sized Arabian filly with angular limb deviation [41], and a 4yo Welsh Pony colt with ventricular septal defect, scoliosis, and facial asymmetry [30]. One of the trisomy 30 EPLs appeared to have prominent cervical curvature, and severe head deformities, while the other lacked protrusion of neck and muzzle, with an unusual positioning of the spine. All four of these trisomy 30 individuals appear to be connected by skeletal abnormalities, suggesting the genetic imbalance associated with trisomy 30 disrupts mechanisms involved with correct skeletal development.

The phenotype of the identified aneuploid EPLs was variable with gestational ages (ranging from 14 to 64 days), fetal and clinical presentations. Contrary to our initial hypothesis, aneuploidy EPLs were not significantly associated with an anembryonic status with only 1/12 aneuploidy pregnancies phenotypically presenting as anembryonic and the proportion of anembryonic aneuploidy EPLs not differing to the proportion of anembryonic diploid EPLs. Based on the variables we studied, an aneuploidy pregnancy was also clinically indistinguishable from diploid pregnancies. Unfortunately, due to the low numbers, we were unable to investigate individual aneuploidy subtypes to identify clinical features associated with a particular aneuploidy type. However, it is of note that we found that mares who suffered an aneuploid EPL, were significantly more likely to carry the next pregnancy to term, compared to those who lost a diploid pregnancy (after removing mares who lost after endometrial cup formation and those who were intentionally not recovered). This suggests that aneuploidy events may occur spontaneously and sporadically in some mares, and further, echoing aneuploidy miscarriage which appears to have a slightly beneficial effect for women, in terms of their chances of having a live birth from the subsequent pregnancy [67].

In conclusion, our results provide the first evidence of aneuploidy in naturally occurring equine pregnancy losses, adding to the overwhelming evidence that the mare provides a far superior system for studying the mechanisms of aneuploidy, compared with rodents. We also support the use of SNP arrays for high throughput identification of aneuploidies in other domesticated species, at a lower per sample cost than conventional technologies.

## Materials and methods

### Ethics statement

All conceptus recoveries from clinical cases of pregnancy loss were performed with owner consent under ethics approval from the Clinical Research and Ethical Review Board at the Royal Veterinary College (URN:2012-1169 and URN:2017-1660-3). Animal care and conceptus recoveries from the clinically normal pregnancies were performed in accordance with the Animals (Scientific Procedures) Act 1986 guidelines set by the Home Office and Ethics Committee of the Royal Veterinary College, London (HO licence PPL 70/8577).

### Sample acquisition, handling and phenotyping

Conceptus material from 55 early pregnancy losses (EPLs) were submitted from veterinary practices across the UK and Ireland, over the 2013-2018 breeding seasons. Pregnancies, confirmed by transrectal ultrasound at 14-16 days post ovulation, and subsequently lost before 65 days post ovulation were included in this study. Following confirmation of pregnancy failure (no heartbeat/collapsed vesicle), conceptuses were recovered by non-invasive uterine lavage by the attending veterinary surgeon according to previously established protocols [38] before being placed into sterile transport media medium (Hank’s Balanced Salt Solution (Sigma-Aldrich, UK), 5% FBS (ThermoFisher, UK), Amphotericin B 250 μg/ml (Sigma-Aldrich, UK), Penicillin-Streptomycin Solution 10000 units/ml (Penicillin), 10000 μg/ml (Streptomycin) (ThermoFisher (Invitrogen), UK), Kanamycin (ThermoFisher (Gibco), UK)), and transported to the Royal Veterinary College (RVC) for assessment and dissection. Reproductive histories of the mares were obtained from the submitting veterinary practices and stud farms, including the mare’s age, records of prior pregnancy losses, history of endometrial pathologies, and previous genetic investigations.

Control conceptuses were obtained from manually terminated clinically normal pregnancies (CNP). Two of the CNPs came from pregnant Thoroughbred broodmares euthanised for complications unrelated to pregnancy. Five Thoroughbred mares aged between 2 and 20 years were housed at the Biological Services Unit (BSU) of the RVC (kept on grass *ad lib* and supplemented with hay over the winter) were donors for the remaining eight CNPs recovered during the 2018 breeding season. Each mare tested negative for Equine Viral Arteritis (EVA) / Equine Infectious Anaemia (EIA), *Taylorella equigenitalis*, *Pseudomonas aeruginosa*, and *Klebsiella pneumonia* (Rossdales Laboratories, Newmarket, UK) prior to the onset of the stud season. Transrectal palpation and ultrasonography identified signs of oestrus and an endometrial swab was submitted for bacteriological culture. Ovulation was induced with 1500 IU of human chorion gonadotropin (hCG, Chorulon; MSD Animal Health, UK) administered intravenously and mares were artificially inseminated 24 hours later with a commercial dose of chilled semen from Thoroughbred stallions of proven fertility. Pregnancies were confirmed 14 days post ovulation by transrectal ultrasound and the development of conceptuses was followed by subsequent ultrasound examinations twice weekly. Clinically normal developing pregnancies (presence of corpus luteum, absence of intrauterine fluid, appropriately sized embryonic vesicle, detection of embryo proper, and appropriately timed detection of fetal heartbeat) were manually terminated between 29 – 41 days gestation and recovered, as previously described [68]. EPL and CNP conceptuses were washed three times in PBS (containing 10 units/ml Penicillin and 10 μg/ml Streptomycin; ThermoFisher (Invitrogen), UK) before the tissues were identified and dissected. Photographs were taken of the different tissues, and seven gross features noted. Tissue samples harvested depended on the developmental age of the conceptus and completeness of the submitted membranes. When available, they included chorion, allantochorion, yolk sac, chorionic girdle, and fetus.

Placenta from 5 healthy term births (2017 stud season) were collected from stud farms in Hertfordshire and Suffolk and brought to the RVC for dissection. Sections were washed three times in PBS and then snap frozen in liquid nitrogen before storage at −80°C. Whole blood from five reproductively normal mares (dams of CNP conceptuses) was collected in heparin vials and peripheral blood mononuclear cells (PBMCs) were isolated as previously described [69]. PBMCs were then snap frozen in liquid nitrogen and transferred to −80°C for long term storage.

### DNA extraction, and sexing of conceptuses

DNA from placental (allantochorion and chorion) and fetal (hind limb and/or tail) tissues (approx. 5×5mm sections, stored at −80°C) and PBMCs were extracted using QIAGEN DNeasy Blood and Tissue kit (Qiagen Sciences, Maryland, USA), following manufacturer’s guidelines. Briefly, tissue or cells were incubated at 56°C overnight in buffer ATL and proteinase K. Tissues were incubated at room temperature for 2 minutes with 28 U RNase A as recommended by the manufacturer then passed through a spin column, before elution with 100 μl Buffer AE provided in the kit. DNA was quantified using a DeNovix Spectrophotometer (DeNovix, Delaware, USA).

Conceptuses were sexed by standard PCR as previously reported [70]. Briefly, primers (Eurofins Genomics, Ebersberg, Germany) for the Sex determining Region of Y (*SRY*) were used to determine the presence (male) or absence (female) of *SRY* from 50 ng of total genomic DNA (S1 Fig) using FIREPol® DNA polymerase (Solis Biodyne, Estonia). Primers for the X-linked Androgen Receptor *(AR*) gene were used as positive control for DNA quality.

Genomic DNA from confirmed male and female equines served as positive controls, while ddH_2_O was used as a negative control. PCR conditions were as follows: 95°C for 5 minutes, 40 cycles of: 95°C for 30 seconds, 60°C for 40 seconds, and 72°C for 1 minute, followed by 72°C for 10 minutes. Amplicons were resolved on a 2% agarose gel and visualised in ultraviolet light.

### HD 670k Equine SNP Array

The 55 EPL samples genotyped on the Axiom™ Equine 670K SNP Genotyping Array [71] included 24 male and 51 female EPL conceptuses from Thoroughbred, Warmblood, and unknown breed pregnancies, with a spread of mare ages and gestational ages. Mare ages ranged from 3 to 21 years old. We also genotyped samples from 10 individual CNP conceptuses, all from Thoroughbreds (both males and females represented), 5 healthy term placentae, all from Thoroughbred pregnancies, and 5 reproductively sound adult Thoroughbred mares (age range 2-20 years). Of the 65 EPL/CNP conceptuses, 14 had matching fetal and placental DNA genotyped by the array. Isolated DNA was submitted to Neogen Europe, Ltd (Auchincruive, Scotland) under a service contract, in which equal loads of DNA (760 ng) were hybridised along with fluorescent probes (PE at 660nm and FAM at 578nm) to the array. One conceptus was represented 5 times on the array to act as both tissue (three individual sections of allantochorion; ALC) and technical (same DNA aliquot represented three times) replicates.

Raw intensity .CEL files were imported in Axiom Analysis Suite (ThermoFisher, UK). Following the CNV Discovery workflow, files were generated that were imported into Integrated Genome Viewer (IGV; [72]) along with EquCab3.0 reference genome. Visualisation of the whole genome at the chromosome level allowed for identification of aneuploidies, along with large structural abnormalities. Quality control metrics were in place using Axiom Analysis Suite, with 86.46% pass rate for the samples.

### Whole Genome Sequencing

Twelve samples (n=7 EPL, n=5 CNP) underwent whole genome sequencing (WGS) using Illumina NovaSeq 6000 at 30X (150 bp pair-end) coverage then aligned to EquCab3.0 reference genome, before applying GATK Best Practices Workflow [73]. EPL samples selected for WGS had a phenotype of Thoroughbred, gestation age between 30 and 42 days, negative uterine swab at loss, with equal representation of conceptus sex and maternal age. CNP conceptuses were age and sex matched to EPL samples. Whole chromosome copy number was determined using SAMtools [74] genome coverage function.

### Digital droplet PCR

Digital droplet PCR (ddPCR) was performed using a BioRad QX200 system (BioRad, Watford, UK). Primers (Eurofins Genomics) were designed (using Primer3Plus; [75]) for two genes per chromosome of interest: *NRG1* and *ANGPT2* (chromosome 27), *SHTN1* and *ACTC1* (chromosome 1), with *MCM6* (chromosome 18) acting as the reference as all individuals were diploid for this chromosome (S1 Fig). Copy number was determined in duplicate in reactions containing 50 ng DNA, using EvaGreen chemistry (final concentration: 100nM each primer, 1x ddPCR Supermix for EvaGreen). C1000 Touch Thermal Cycler performed ddPCR reactions as follows: 95°C for 5 minutes, 40 cycles of: 95°C for 30 seconds and 58°C for 1 minute, 4°C for 5 minutes, and 90°C for 5 minutes. Droplets were analysed using Bio-Rad QX200 Droplet Reader and QuantaSoft software (BioRad). The copy number of each product was manually calculated relative to *MCM6*.

### Statistical Analysis

Statistical analysis for numerical variables was performed using GraphPad Prism 7.03 (GraphPad Software, California, USA). Normality of distributions for each group were assessed using Shapiro-Wilk normality test. Where both groups passed the normality test, a two-tailed T-Test was performed. Where one or more groups did not pass normality tests, a Mann-Whitney test was applied. Significance was set at p<0.05 for both statistical tests. Simple linear regression was calculated using GraphPad Prism 7.03. Factors that were determined prior to the pregnancy loss (including the sex of the conceptus, the mare status at the beginning of the season, mare and stallion age, breed, use of ovulation induction, twin pregnancy, and uterine infection) were analysed to identify variables that differentiated aneuploid EPLs from non-aneuploid EPLs. Due to the low numbers, barren (did not produce a live foal from previous season) and rested (deliberately not bred) mares were combined into one group and compared with maiden (never bred) and foaled (produced a live foal from previous season) mares. Other variables investigated occurred after the loss (whether she ended the season with a foal) and aimed to identify whether an aneuploidy EPL at the beginning of the season would decrease her chance of a successful season. The categorical variables were analysed in SPSS (v26), using Fisher’s exact test (significance set at 0.05). As only one variable was significant, a multivariable linear regression was deemed unnecessary.

## Acknowledgements

The authors would like to thank all the participating stud farms and veterinarians for their continued support. Thanks also to Dr Daniel Hampshire and Dr Belinda Rose for assistance processing the conceptuses, and Dr Ruby Chang for statistical advice.

## Author contribution

AMdM, TR, and DCW, were involved in the conceptualisation of the project and acquisition of funding. TR provided the PCR protocol to sex the conceptuses. CAS performed the formal analysis, and along with AMdM prepared the original manuscript draft. BWD, AK, JRC, JC, and AJM provided vital resources in terms of computational resources, and animal samples. AK, AMdM, JRC, JC, and AJM provided invaluable expert knowledge for the interpretation of results generated in the clinical analysis. AMdM, DCW, TR, and AK provided extensive comments and edits of the original manuscript draft. TR, BWD oversaw the supervision of the analysis of the SNP array and WGS methodologies, while AMdM oversaw the supervision of other methodologies and the wider research activity.

## Supporting Information

**S1 Table. Details of primers used for both standard and digital droplet PCR (ddPCR)**.

**S2 Table. Analysis of 10 clinically relevant variables**. No variables measured could distinguish between and aneuploid loss and a euploid loss

